# Activation of lateral hypothalamic group III mGluRs suppresses drug-seeking following abstinence and cocaine-associated increases in excitatory drive to orexin/hypocretin cells

**DOI:** 10.1101/360081

**Authors:** Jiann W. Yeoh, Morgan H. James, Cameron D. Adams, Jaideep S. Bains, Takeshi Sakurai, Gary Aston-Jones, Brett A. Graham, Christopher V. Dayas

## Abstract

The perifornical/lateral hypothalamic area (LHA) orexin (hypocretin) system is involved in drug-seeking behavior elicited by drug-associated stimuli. Cocaine exposure is associated with presynaptic plasticity at LHA orexin cells such that excitatory input to orexin cells is enhanced, both acutely and into withdrawal. These changes may augment orexin cell reactivity to drug-related cues during abstinence and contribute to relapse-like behavior. Studies in hypothalamic slices from drug-naïve animals indicate that agonism of group III metabotropic glutamate receptors (mGluRs) reduces presynaptic glutamate release onto orexin cells. Therefore, we examined the group III mGluR system as a potential target to reduce orexin cell excitability *in*-*vivo*, and tested whether activating these receptors could normalize orexin cell activity following cocaine and reduce cocaine-seeking elicited by drug-associated stimuli during abstinence. First, we verified that group III mGluRs regulate orexin cell activity *in vivo* by showing that intra-LHA infusions of the selective agonist L-(+)-2-Amino-4-phosphonobutyric acid (L-AP4) reduces Fos expression in orexin cells following 24h food deprivation. Next, we extended these findings to show that intra-LHA L-AP4 infusions reduced discriminative stimulus-driven cocaine-seeking following withdrawal. L-AP4 had no effect on general motor activity of sucrose self-administration. Finally, using whole-cell patch clamp recordings from identified orexin cells in orexin-GFP transgenic mice, we show that enhanced presynaptic drive to orexin cells persists for up to 14d into withdrawal and that this plasticity is normalized by L-AP4. L-AP4 had no effect on measures of postsynaptic plasticity in cocaine-exposed animals. Together, these data indicate that agonism of LHA group III mGluRs reduces orexin cell activity *in*-*vivo* and is an effective strategy to suppress cocaine-seeking behavior following withdrawal. These effects are likely mediated, at least in part, by normalization of presynaptic plasticity at orexin cells that occurs as a result of cocaine exposure.

## 1. INTRODUCTION

Orexin (hypocretin) cells in the perifornical/lateral hypothalamic area (LHA) play a key role in motivated behavior, including drug-seeking ((Harris *et al.*, 2005; Mahler *et al.*, 2014; Sakurai, 2014; James *et al.*, 2017b). The orexin system is particularly important for driving drug-seeking behavior elicited by environmental cues associated with drug use (for review, see James *et al.*, 2017). For example, orexin cells are recruited by discriminative and contextual stimuli previously paired with drug (Dayas *et al.*, 2008; Moorman *et al.*, 2016; Martin-Fardon *et al.*, 2018), and the magnitude of orexin cell activation is strongly correlated with drug-seeking behavior (Harris *et al.*, 2005; Richardson & Aston-Jones, 2012; Moorman *et al.*, 2016). Furthermore, orexin receptor antagonists, particularly those acting at the orexin-1 receptor, are highly effective at blocking stimulus-driven drug seeking across various drugs of abuse (Smith *et al.*, 2009; Smith *et al.*, 2010; Plaza-Zabala *et al.*, 2013; Martin-Fardon & Weiss, 2014; Moorman *et al.*, 2017), but these compounds have no effect on drug-seeking elicited by drug priming injections (Zhou *et al.*, 2012). Thus, the orexin system offers a promising target for therapeutic interventions designed to ameliorate craving in response to drug-associated stimuli during abstinence (Yeoh *et al.*, 2014a; James *et al.*, 2017b).

Excitatory synapses onto orexin cells rapidly rewire in response to environmental and physiological challenges, including food restriction, sleep deprivation and drug exposure/withdrawal (Horvath & Gao, 2005; Yeoh *et al.*, 2012; Rao *et al.*, 2013). We and others have reported that cocaine exposure enhances excitatory drive to LHA orexin cells and that these changes persist into acute withdrawal (Yeoh *et al.*, 2012; Rao *et al.*, 2013). Such changes may drive enhanced orexin output in response to drug-associated cues during abstinence and contribute to relapse-like behavior. Thus, suppression of excitatory drive to LHA orexin circuits during withdrawal might represent a novel strategy to reduce relapse risk (Yeoh *et al.*, 2012). To this end, previous work has shown that metabotropic glutamate receptors (mGluRs) in hypothalamus maintain tonic presynaptic inhibition at excitatory (and inhibitory) synapses (Kuzmiski *et al.*, 2009; Kuzmiski & Bains, 2010). In particular, the group III mGluR subtype has been shown to gate presynaptic glutamate release onto orexin cells, such that activation of these receptors with the selective agonist L-(+)-2-Amino-4-phosphonobutyric acid (L-AP4) reduces presynaptic glutamate release onto orexin cells *in vitro* (Acuna-Goycolea et al., 2004). Currently, it is unclear whether agonism of group III mGluRs reduces orexin cell activity *in vivo*, or whether this approach could be used to suppress enhanced excitatory input to orexin cells following cocaine.

Here, we sought to examine the effect of infusing L-AP4 directly into LHA in freely behaving rats, both in terms of orexin cell activity and drug-seeking behavior. First, we evoked Fos in orexin cells using food deprivation, a stimulus known to enhance excitatory drive to orexin cells (Horvath & Gao, 2005), and show that Fos is suppressed by intra-LHA L-AP4 infusions. In rats with a history of cocaine self-administration, we show that intra-LHA L-AP4 injections reduces drug-seeking elicited by discriminative stimuli following 14d withdrawal. Next, we sought to verify LAP-4’s mechanism of action by carrying out whole-cell patch clamp recordings in orexin-GFP mice with a history of cocaine exposure. We extend upon previous findings to show that the 14d withdrawal time point is associated with enhanced measures of presynaptic and postsynaptic plasticity to orexin cells, and that L-AP4 selectively normalizes withdrawal-induced increases in excitatory drive. Together, our findings highlight the group III mGluR system as a potential mechanism through which drug-induced plasticity to orexin cells can be normalized, leading to reduced drug-seeking elicited by drug-associated stimuli during abstinence.

## 2. MATERIALS & METHODS

### 2.1 Experiment 1: Behavioral studies in rats

#### 2.1.1 Animals

Rats weighing 200-250g upon arrival (Charles River,USA) were single-housed on a reversed 12hr light/dark cycle (lights off at 0800) with food and water available *ad libitum*. All procedures were carried out in accordance with the Institutional Animal Care and Use Committee of Rutgers University. All experiments were carried out in the animals’ active period.

#### 2.1.2. Surgery for intravenous catheter implantation

Rats were handled daily for 1wk before undergoing intravenous (i.v) catheter surgery as described previously (James *et al.*, 2016; James *et al.*, 2018). Briefly, animals were anaesthetized with isoflurane (1-3Q% with a flow rate of 2L/min) and received the analagesic rimadyl (5mg/kg, s.c.). Chronic indwelling catheters were inserted into the right jugular vein and exited the body via a port between the scapulae. Rats received prophylactic i.v. cefazolin (10mg) and heparin (10U) daily starting 1 day after surgery and continuing throughout self-administration training.

#### 2.1.3. Surgery for LH-directed cannulae

Animals were placed in a stereotaxic frame, anaesthetized as above, and underwent stereotaxic surgery for guide cannulae (23G;PlasticsOne) implantation 2mm above the fornix (AP:-2.65; ML:±1.90; DV:-7.25). Cannulae were affixed to the skull with four stainless steel screws and dental cement, and were occluded with 26G gauge bilateral steel stylets (PlasticsOne).

#### 2.1.4. Food deprivation and Fos expression experiments

A subset of rats (n=5) previously used for sucrose self-administration and locomotor testing (described below) were used in this experiment. These rats were maintained on an *ad libitum* diet for all other tests; here they were subjected to 24h food deprivation, which is known to enhance mEPSCs frequency and induce c-*fos* expression in orexin cells (Mieda *et al.*, 2004; Horvath & Gao, 2005). Rats received unilateral microinjections of L-AP4 (100nmol) and aCSF into opposite hemispheres (hemispheres counterbalanced across animals). This dose of L-AP4 was selected based on previous studies indicating that injections of this dose into striatum blocks cocaine hyperlocomotion without affecting general motor behavior (Mao & Wang, 2000). Ninety minutes later, animals were anesthetized (ketamine/xylazine) and then transcardially perfused with saline followed by 4% paraformaldehyde (PFA). Brains were removed and post-fixed overnight, and then transferred to a 20% sucrose solution for storage until they were sectioned into 40um sections on a cryostat.

Tissue from LHA was processed for Fos and orexin using procedures outlined elsewhere (Yeoh *et al.*, 2012). Briefly, tissue underwent three washes in PBST-azide (phosphate buffer saline + 0.03% Triton X-100 + azide), before being incubated overnight at room temperature in primary antibodies goat anti-orexin-A (1:500; Santa-Cruz Biotechnology, catalog number SC-8070; validation details at JCN Antibody database AB_653610) and rabbit anti-Fos (1:1000; Synaptic Systems, catalog number 226003; validation details at JCN Antibody database AB_2231974) in 2% normal donkey serum. The next day, the tissue was washed three times in PBST and incubated for 2h in appropriate secondary antibodies coupled to Alexa-Fluor 488/594-conjugated donkey anti-goat/rabbit (Jackson Immunoresearch Laboratories). Sections were then rinsed in PB, mounted onto glass slides and cover slipped using Fluoroshield mounting medium with DAPI (Abcam). Hypothalamic sections were imaged in tiles with a 16x objective on a Zeiss AxioZoom V16 microscope and then stitched together using Zen Imaging software. Cell counts were also performed using the Zen Imaging software. The number of cells immunoreactive for orexin-A, as well as the number of orexin-A cells that also expressed Fos, were quantified in three sections taken through main rostrocaudal extent of the hypothalamic orexin cell region in each rat by an experimenter blind to hemispheric treatment conditions. Counts were carried out in a 300×300μ m region immediately ventral to the injector tip in each hemisphere. The percentage of orexin-immunoreactive cells that co-expressed Fos was averaged across each hemisphere for each subject, then compared across L-AP4 versus aCSF microinjected hemispheres across all subjects.

#### 2.1.5. Cocaine self-administration training

A separate group of rats (n=16) were trained to self-administer cocaine in standard Med-Associates operant chambers. Each chamber was equipped with two retractable levers with white lights above it, a red house-light located at the top of the chamber wall opposing the levers and a tone generator. A syringe pump located outside of the sound-attenuating chamber delivered cocaine intravenously. Data acquisition and behavioral testing equipment were controlled by MED-PC IV program (Med Associates,USA). Seven days after surgery, animals were trained to respond for i.v cocaine hydrochloride infusions. Animals were first trained on a fixed ratio 1 (FR1) schedule of reinforcement for 2hr/day, 6-7days/week. During this period, responding on the active (right) lever resulted in a 3.6s infusion of cocaine (0.2mg/50ul infusion) via the i.v catheter, which was paired with the activation of white cue light above the active lever. A 20s timeout period followed each infusion delivery, which was signaled by the house light turning off, during which additional presses on the active lever were of no consequence. Responses on the inactive (left) lever were recorded but had no scheduled consequences at any time. Rats underwent daily 2hr cocaine self-administration sessions until they met a criterion of >10 cocaine infusions/day over three consecutive days. At this time, rats were transitioned to daily 2hr randomized conditioning sessions for cocaine or saline infusions (FR1), in the presence of distinct discriminative stimuli (‘first round’ of conditioning). We used a discriminative stimulus paradigm, as this form of stimulus is associated with robust activation of orexin cells (Dayas *et al.*, 2008; Moorman *et al.*, 2016; Martin-Fardon *et al.*, 2018). For cocaine (DS+), this involved black-striped wallpaper on the chamber door and a vanilla odor placed in a receptacle under the active lever. The time out period following an active lever response was signaled with the illumination of house light. For saline (DS-), chambers were equipped with black-spotted wallpaper on the chamber door and a lemon odor. Time out periods were signaled by a constant 78dB 2.9kHz tone. After 16d of conditioning (8DS^+^/8DS^−^ sessions, randomized order), animals underwent 13d of abstinence in their home cage where they were left undisturbed except for weekly bedding changes.

#### 2.1.6. Test of drug-seeking after abstinence

Following abstinence animals were returned to the operant chambers, presented with the DS-cues, and tested for drug-seeking behavior for 30min. Immediately following testing, rats were gently restrained and injectors were lowered into the bilateral cannulae (28G, PlasticsOne) and then removed; no injection was made. The following day (14d following final conditioning session), animals underwent testing of drug-seeking following abstinence under DS+ conditions for 30min. 10min prior to testing, animals received a bilateral microinfusion of L-AP4 (100nmol; (Mao & Wang, 2000)) or vehicle (aCSF) into the orexin field (0.5ul over 1min). Injectors were left in place for an additional 1min to limit upward diffusion of the injectate. Rats were then placed in the self-administration chambers under DS+ conditions and tested for drug-seeking behavior for 30 min. The following day, animals were returned to self-administration conditioning training (8DS+/8DS-; ‘second round’ of conditioning), underwent 14d abstinence, and prior to testing for drug-seeking in the DS+ context, received the opposite treatment (L-AP4 or aCSF) to their first test. The order in which animals received the two different treatments was counterbalanced for the group.

#### 2.1.7. Sucrose pellet self-administration

Drug-naïve animals (n=7) were trained to self-administer sucrose pellets (45mg, Test Diet) on an FR1 schedule during 2h sessions (McGlinchey *et al.*, 2016). As with cocaine training, each rewarded active lever response was paired with a light and tone cue, followed by a 20s timeout period. Training continued until the number of active lever presses and pellets earned was stable (<20% variability) over 3 consecutive days. The effect of intra-LH L-AP4 or aCSF infusions on sucrose self-administration was tested under a FR1 schedule for 2h, using injection procedures identical to those used for cocaine (described above). Test days were separated by at least 2d of further FR1 training (<20% variability relative to pre-test baseline).

#### 2.1.8. Locomotor assay

The same rats tested for sucrose self-administration (n=7) were also assessed for general locomotor activity. Procedures are published elsewhere (McGlinchey *et al.*, 2016; James *et al.*, 2018). Briefly, rats were placed in locomotor chambers (clear acrylic, 42cm x 42cm x 30cm) equipped with SuperFlex monitors (Omintech Electronics Inc, Columbus, OH) containing a 16 x 16 infrared light beam array for the x/y axis (horizontal activity) and 16 x 16 infrared light beams for the z axis (vertical activity). Activity was recorded by Fusion SuperFlex software. Animals were habituated to the locomotor boxes for 2h/day for at least 3d, and until the average total distance traveled by each rat was within a range of ±25% of the mean of those days. On test days, total distance traveled, as well as horizontal and vertical beam breaks were recorded over a 2h session. Between test days, animals were again habituated to the boxes until stable behavior was again observed over 3 consecutive days. All animals received L-AP4 and aCSF as described above in a counterbalanced order.

#### 2.1.9. Verification of injector placement

For cocaine rats, injector placement was verified by deeply anesthetizing the animals with isoflurane and then lowering injectors into the cannulae. Rats were then decapitated and brains were flash-frozen in 2-methylbutane and stored at −80°C. Brains were sectioned into 40μm sections on a cryostat, slide mounted, and visually inspected under a 40x microscope. All other experiments (sucrose, locomotor and Fos studies) utilized the same animals – injection sites were verified during cell quantification, as outlined above.

#### 2.1.10 Statistical analyses

For the food deprivation experiment, the percentage of orexin-positive cells co-expressing Fos was compared across treatment groups using a paired *t*-test analysis (with the aCSF-injected hemisphere acting as the control for each animal). In cocaine self-administering rats, self-administration values for the final DS+ and DS-sessions were compared using paired samples *t*-tests. Responses during DS-drug-seeking tests sessions were calculated as the average of both sessions and compared to DS+ drug-seeking values following L-AP4 or vehicle treatment using repeated measures ANOVA. Two animals were excluded from the drug-seeking analyses as they failed to exhibit robust DS+/induced-behavior following vehicle treatment. For these two animals, responding on the DS+ test was lower than their responding on the DS-test and was more than 3SDs below the overall group average.

### 2.2. Experiment 2: Electrophysiological experiments in mice

#### 2.2.1. Animals

All experimental procedures were approved by the University of Newcastle Animal Care and Ethics Committee and performed in accordance with the NSW Animal Research Act, Australia. Breeding pairs of orexin-GFP mice DBF1 (F1 of DBA2XB6J, (Sakurai *et al.*, 2005)) were transferred from Kanazawa University and backcrossed onto C57BL/6J background for at least nine generations at the Animal Resources Centre, NSW, Australia. Mice aged 4-6 weeks were housed 2-4/cage in temperature- and humidity-controlled conditions on a reverse 12hr daylight cycle (lights off 0700) with *ad libitum* access to food and water. All experiments were carried out during the animals’ dark (active) phase.

#### 2.2.2. Confirmation of GFP-orexin cells in the LHA

This mouse line has been well validated and we observed GFP labeling exclusively in the LHA medial and lateral to the fornix as per previous studies (Figure 6C; Yamanaka *et al.*, 2003a; Yamanaka *et al.*, 2003b).

#### 2.2.3. Drugs

Cocaine Hydrochloride (GlaxoSmithKline,Australia) was dissolved in sterile physiological saline: (15mg/ml) for intraperitoneal (i.p.) injection. Picrotoxin and 6-cyano-7-nitroquinoxaline-2,3-dione, CNQX, were purchased from Sigma-Aldrich (MO,USA). L-(+)-2-Amino-4-phosphonobutyric acid (L-AP4) was purchased from Tocris Bioscience (Bristol,UK).

#### 2.2.4. Behavioral Paradigm

Prior to treatment, animals were conditioned to daily handling and subjected to single-daily sham injections, after which they were placed in an enclosed arena (50cm x 50cm) for 30 min. Animals were then randomly allocated into two treatment groups, either saline (n=10) or cocaine (15mg/kg; i.p. n=8) for seven consecutive days (Yeoh *et al.*, 2012; Yeoh *et al.*, 2014b). Immediately following injections, animals were placed in the same enclosed arenas (30mins) before they were returned to their home cage. In a subset of mice (n=3 saline v.s. n=3 cocaine), locomotor activity (total distance traveled) was assessed using Ethovision software. Brain slice electrophysiology experiments were undertaken 14 days after the last cocaine/saline injection session.

#### 2.2.5. Slice Preparation for electrophysiology experiments

Mice were deeply anaesthesized with ketamine (1ml/kg, i.p) and decapitated. Brains were rapidly removed and immersed in ice-cold oxygenated (95%O_2_, 5%CO_2_) sucrose-substituted artificial cerebrospinal fluid (S-ACSF, containing in mM: 236.2 sucrose, 25NaHCO_3_, 13.6glucose, 2.5KCl, 2.5CaCl_2_, 1NaH_2_PO_4_, 1MgCl_2_). Coronal slices (250μm) containing the LHA were obtained using a vibrating microtome (Campden, England), then transferred to an oxygenated storage chamber containing ACSF (119.4mM NaCl substituted for sucrose) and incubated for 1hr at room temperature prior to recording.

#### 2.2.6. Electrophysiology Recording

Slices were transferred to a recording chamber and continually superfused with oxygenated ACSF, maintained between 32-34°C by a temperature controller (Warner Instruments, USA). Whole-cell recordings were made using Multiclamp 700B amplifier (Molecular Devices,CA), digitized at 10kHz, via an ITC-18 computer interface (Instrutech,NY) and recorded onto a Macintosh computer running Axograph X software. All recordings were restricted to the LHA brain region spanning between Bregma −1.06 mm and −1.70 mm (Paxinos *et al.*, 2001). After obtaining whole-cell recording configuration, series resistance and input resistance were calculated based on the response to a −5mV voltage step from a holding potential of –70mV. These values were monitored at the beginning and end of each recording and data were rejected if values changed by more than 20%. A bipolar stimulating electrode was positioned immediately medial and dorsal to the PF/LHA to stimulate excitatory inputs (0.1ms pulse duration, 1.2x threshold stimulus intensity). Evoked EPSCs (eEPSCs) were recorded with an internal solution containing (in mM): 120Cesium methanesulfonate, 20CsCl, 10HEPES, 4Mg-ATP, 0.3Na_3_-GTP, 0.2EGTA, 10NA-phosphocreatine, 5QX-314 (pH7.3 with CsOH). These recordings assessed both the paired-pulse ratio and AMPA:NMDA ratio of eEPSCs. Cells were voltage clamped at −70mV, in the presence of picrotoxin (100μM, Yamanaka *et al*, 2003) to record the paired-pulse ratio (50ms interstimulus interval) of AMPAR-mediated EPSCs. NMDAR-mediated EPSCs were recorded at a holding potential of +40mV with the addition of CNQX (10μM) to abolish AMPA-receptor mediated currents. Miniature excitatory postsynaptic currents (mEPSCs) were recorded in voltage clamp at a holding potential of −70mV with series resistance of <25MΩ, in the presence of tetrodotoxin (1μM) and picrotoxin (100μM). In addition, L-AP4 (50μM) was bath applied to determine the effects of GPIII mGluR activation. Recording pipettes (6-9MΩ) for mEPSC recordings contained an internal solution containing (in mM): 135KCH_3_SO_4_, 6NaCl, 2MgCl_2_, 10HEPES, 0.1EGTA, 2MgATP, 0.3NaGTP (pH7.3 with KOH) and 0.1% Neurobiotin.

#### 2.2.7. Data Analysis

Peak amplitude, rise-time and decay time constant were evaluated for eEPSCs using semi-automated procedures in the Axograph X software. Paired-pulse ratio (PPR) was calculated by dividing the mean peak amplitude of the second eEPSC by the mean peak amplitude of the first. AMPA:NMDA ratios were calculated by dividing the mean peak amplitude of evoked AMPA EPSCs (recorded at −70mV) by the mean peak amplitude of evoked NMDA EPSCs (recorded in CNQX at +40mV). mEPSCs were detected and captured using a sliding template method (Clements and Bekkers, 1997), along with a minimum amplitude threshold criteria of 10pA. Captured mEPSCs were individually inspected and excluded from the analysis if they included overlapping events or had an unstable baseline before the rise or during the decay phase of the mEPSC. Analyses were performed on averaged mEPSCs, generated by aligning the rising phase of all accepted events. Peak amplitude, rise-time (calculated over 10-90% of peak amplitude) and decay time constant (calculated over 20–80% of the decay phase) were obtained using semi-automated procedures. Average mEPSC frequency was obtained by dividing the number of captured events by the analysis duration in seconds.

#### 2.2.8. Statistical analyses

For sensitization behavior, data were grouped into ‘baseline’ (last 3d of habituation), ‘early’ (first 3d of cocaine/saline treatment) and ‘late’ (last 3d of cocaine/saline treatment), and compared using repeated-measures ANOVA. All electrophysiological data were compared using either Student’s *t*-tests or one-way ANOVA.

## 3. RESULTS

### 3.1. Effects of L-AP4 on orexin cell activity and drug-seeking behavior in-vivo

#### 3.1.1. Microinfusions of L-AP4 into the LHA reduced food-deprivation-induced Fos-protein in orexin cells

To test whether intra-LHA injections of L-AP4 can alter orexin cell activity *in vivo*, we infused L-AP4 and aCSF unilaterally in opposite hemispheres following 24h food deprivation. The total number of orexin-immunoreactive cells quantified was similar across aCSF- and L-AP4-treated hemispheres (*P*>0.05, Figure 1A). Consistent with previous studies (Lutter et al, 2008; Mieda et al, 2004), approximately 75% of orexin-immunoreactive cells co-expressed Fos following food deprivation in the aCSF control group. This percentage was significantly lower (~40%) in the L-AP4-treated group (t_4_=9.774, *P*=0.0006, paired-samples t-test; Figure 1B, 1C), indicating that activation of group III mGluRs with L-AP4 suppressed orexin cell activity.

**Figure 1.**
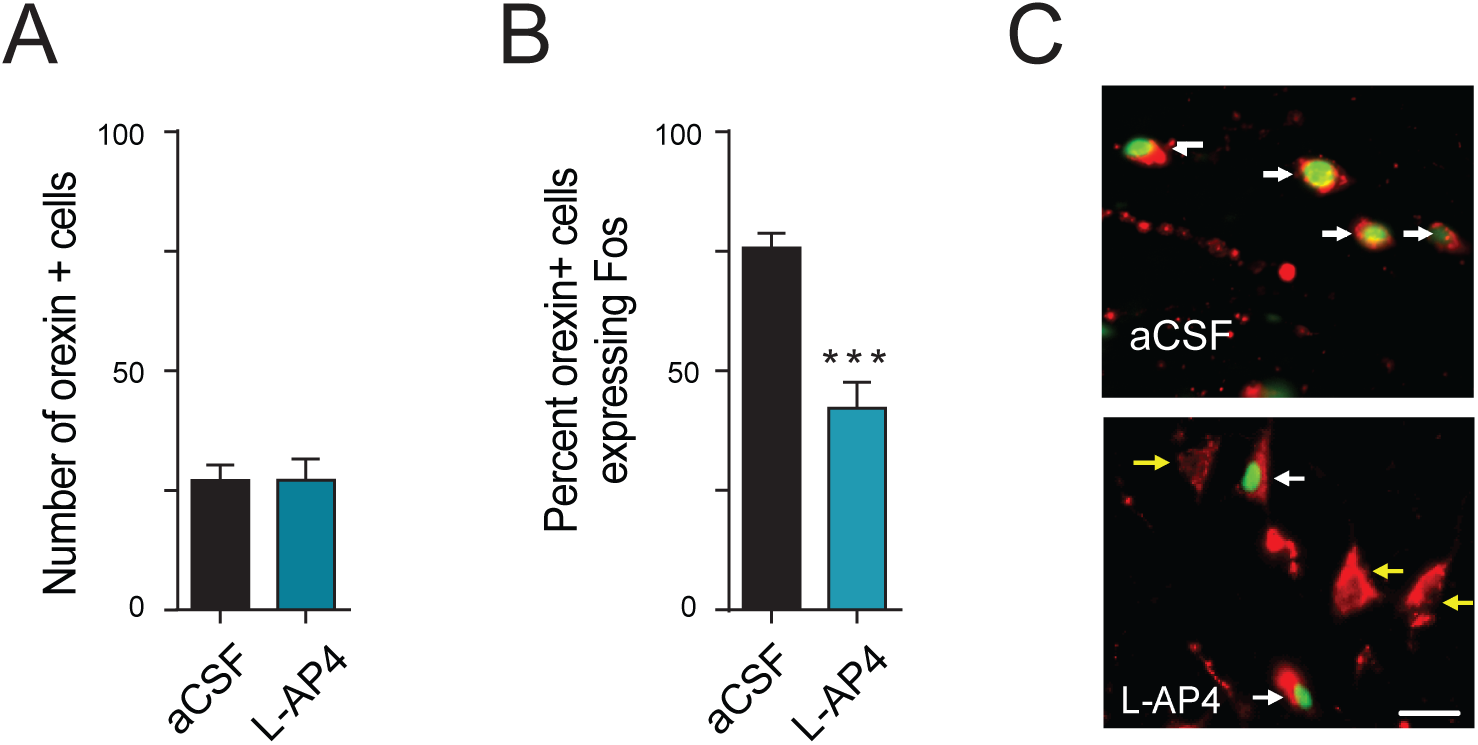
Intra-LHA microinfusions of L-AP4 attenuate orexin cell activity. Activation of Fos-expressing orexin cells was quantified in food-deprived rats following intra-LH infusions of L-AP4 into one hemisphere, and aCSF into the opposite hemisphere. *A*, The total number of orexin-immunoreactive cells quantified in L-AP4-injected hemispheres was identical to that of the aCSF-injected hemispheres. *B*, L-AP4 microinfusions into LHA significantly reduced the number of orexin-immunoreactive cells that co-expressed Fos (^***^*P*<0.001). *C*, Representative photomicrographs from the same animal showing Fos (green) in orexin (red) cells following intra-LHA injections of aCSF (top panel) and L-AP4 (bottom panel) in opposing hemispheres. Scale bar = 25μm. White arrows indicate cells immunoreactive for orexin and Fos. Yellow arrows indicate cells immunoreactive for orexin only.

#### 3.1.2. Microinfusions of L-AP4 into LHA reduced cocaine-seeking following 14d abstinence

Animals met criterion for stable responding during initial FR1 training in an average of 4.63 (±0.70 SEM) days. Animals learned to discriminate between DS+ and DS-stimuli during the first round of conditioning, as evidenced by a significantly greater number of active lever presses (t_15_=4.246, *P*=0.0007) and infusions (t_15_=7.202, *P*<0.0001) on the final DS+ session compared DS-session (Figure 2A). Inactive lever responding was significantly higher on the final DS-sessions compared to the DS+ sessions (t_15_=6.146, *P*<0.0001; Figure 2A). The number of active and inactive lever responses, as well as the number of infusions earned, during the second round of conditioning were almost identical to those in the first round (*P*’s>0.05; data not shown). There was no difference in self-administration behavior between L-AP4 and aCSF-treated animals on the second round of conditioning (*P’*s>0.05).

**Figure 2.**
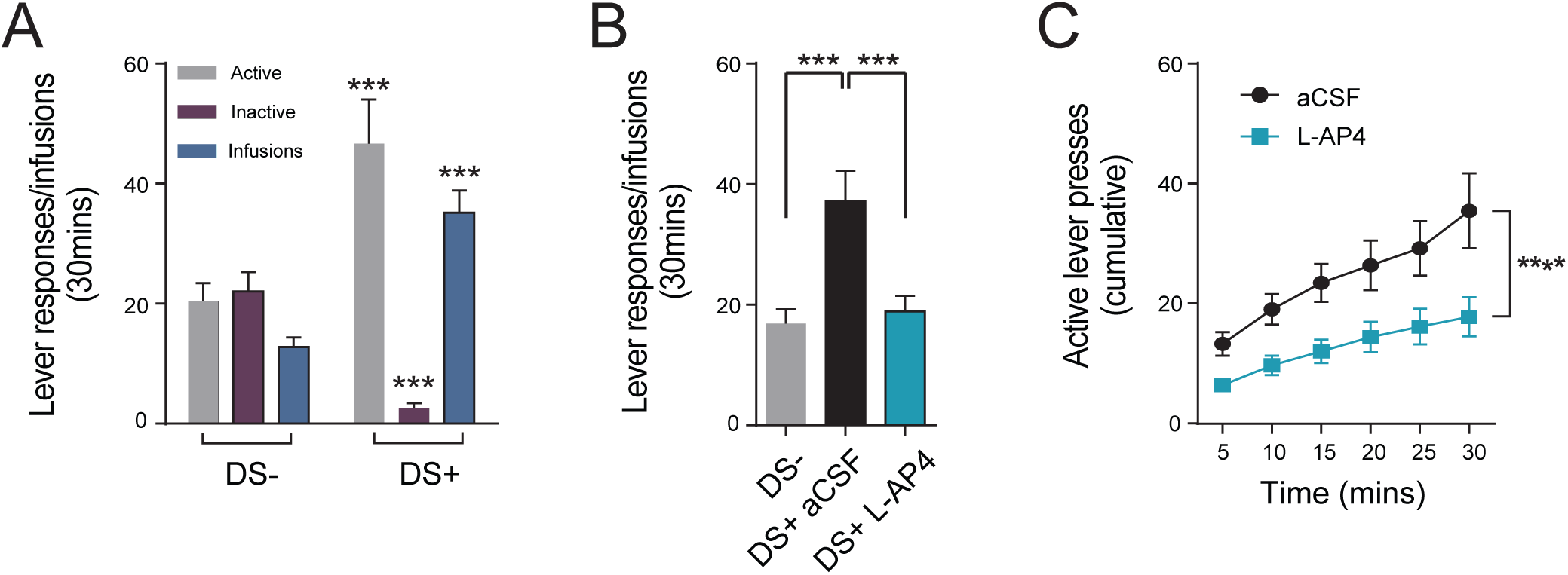
Intra-LHA microinfusions of L-AP4 attenuate cocaine seeking behavior elicited by drug-associated discriminative stimuli following 14d abstinence. *A*, Rats were trained to self-administer cocaine (DS+) or saline (DS-) in the presence of discriminative stimuli. Rats learned to discriminate between DS+/DS-stimuli, as indicated by higher active lever responses and infusions during the final DS+ session, relative to the final DS-session of the first round of conditioning (^***^*P*<.001). Behavior during the second round of conditioning was indistinguishable from the first (data not shown). *B*, Following 14d withdrawal, testing (30min) under DS+ conditions following was associated with significantly higher levels of active lever responding compared to testing under DS-conditions in aCSF-treated rats. Active lever responding in the DS+ test was significantly attenuated following intra-LH microinfusions of L-AP4 (^***^*P*<0.01). *C*, Comparison of cumulative lever responding across the DS+ test indicates that lever pressing was significantly attenuated by L-AP4 across the entire duration of the test session (^****^*P*<0.0001).

To test the effect of group III agonism on drug-seeking behavior, we infused L-AP4 or vehicle directly into LHA immediately prior to testing animals for drug-seeking in the presence of DS+ stimuli following 14d homecage abstinence. Re-exposure to DS+ stimuli following aCSF microinfusions was associated with significantly greater levels of drug-seeking (active lever responding) compared to testing under DS-conditions (F_2,32_=10.70, P=0.0042; Holm-Sidak post-hoc comparison, *P*=0.0014; Figure 2B). This DS+-elicited drug-seeking was significantly attenuated by intra-LHA infusions of L-AP4 (*P*=0.0010, Holm-Sidak post-hoc test; Figure 2B). LAP-4-induced suppression of drug-seeking was evident across the entire duration of the test session (main effect of ‘treatment’, F_5,120_=37.68, P<0.0001, two-way ANOVA; Figure 2C). There was no effect of L-AP4 on inactive lever responding during the DS+ test (*P*>0.05, data not shown).

#### 3.1.3. Microinfusions of L-AP4 into LHA had no effect on sucrose self-administration behavior or general locomotor activity

To test for any non-specific effects of intra-LHA L-AP4 infusions, we made bilateral injections of the group III mGluR agonist prior to testing for sucrose self-administration and spontaneous locomotor activity. Animals achieved stable sucrose self-administration behavior in 6.87±0.93(SEM) days. On test day, infusions of either aCSF or L-AP4 did not affect the number of responses made on the active across the 30min test session (*P*>0.05; Figure 3A,B), nor the number of sucrose pellets earned during the session (*P*>0.05; Figure 3C).

**Figure 3.**
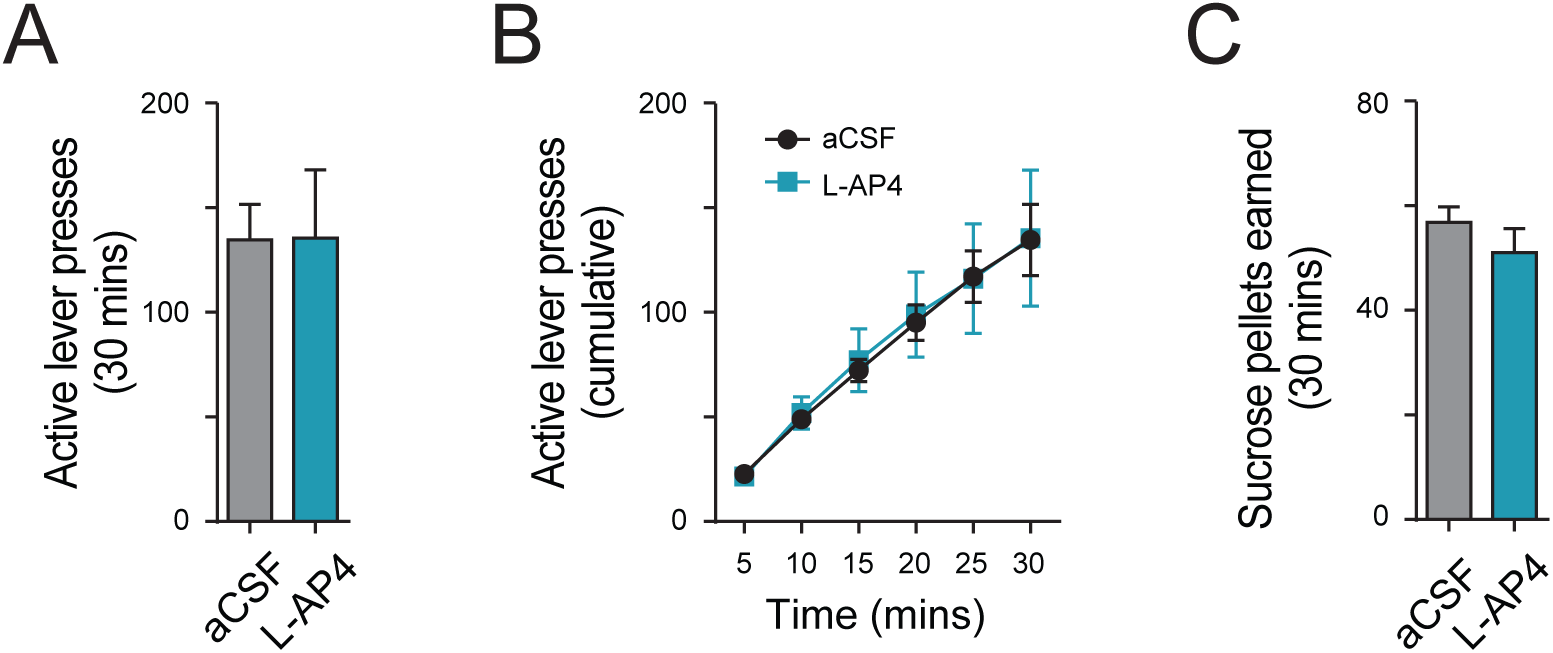
LHA microinfusions of L-AP4 have no effect on high-rate responding for sucrose pellets. We tested whether intra-LH L-AP4 infusions affected general reward seeking, as assessed by responding for sucrose on an FR1 schedule. *A*, L-AP4 had no effect on the total number of active lever responses made in the 30min test session. *B*, Active lever responding for sucrose was similar between L-AP4- and aCSF-treated animals at all time points during the test. *C*, L-AP4 did not affect the total number of sucrose pellets earned during the 30min test session.

Similarly, there was no effect of treatment on total distance travelled, (Time x Treatment interaction, *P*>0.05; Figure 4A,B), horizontal activity (*P*>0.05; Figure 4C) or vertical activity (*P*>0.05; Figure 4D) during locomotor testing, indicating that L-AP4 infusions did not affect general motoric activity. Interestingly, L-AP4 treatment significantly increased time spent in the centre square of the open field during locomotor testing (t_6_=2.96, *P*<0.05; Figure 4E), indicating that L-AP4 reduced anxiety-like behavior during this test.

**Figure 4.**
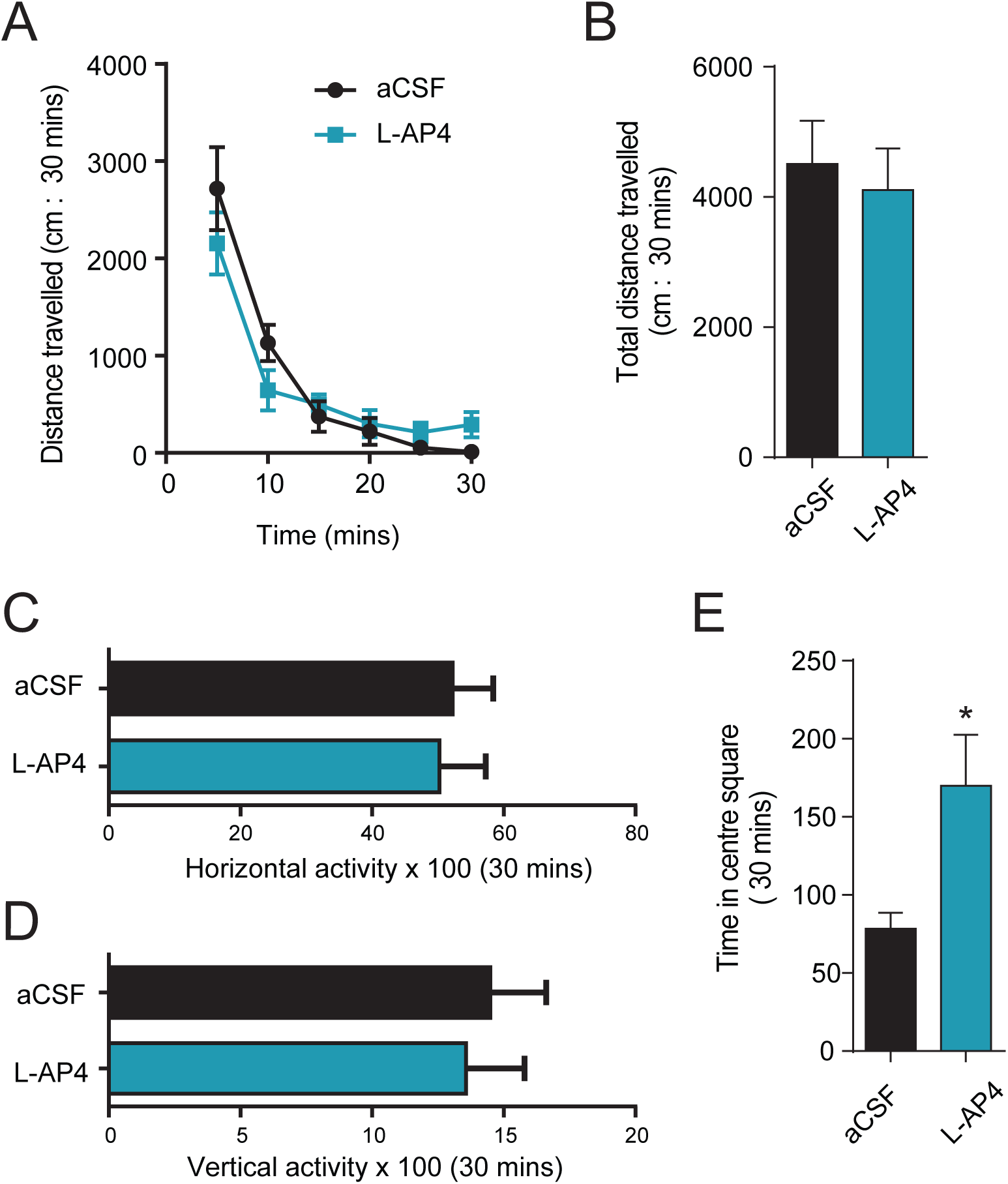
Intra-LHA microinfusions of L-AP4 do not affect general motor behavior. *A*, On a 30min test of general locomotor activity, intra-LHA L-AP4 microinfusions had no effect on the distance traveled in the locomotor test, in any 5min epochs across the 30min test. *B*, Similarly, L-AP4 did not affect the total distance traveled across the entire duration of the 30min test. *C*, L-AP4 did not affect horizontal activity. *D*, L-AP4 did not affect vertical activity. *E*, Consistent with evidence that orexin signaling has anxiogenic properties, intra-LHA microinfusions of L-AP4 significantly increased the amount of time spent in the centre square of the open field during the locomotor test. ^*^*P*<0.05.

#### 3.1.4. Histology

For drug-seeking experiments, microinjections (L-AP4 and vehicle) made into the LHA ranged from 2.64 mm to 3.12 mm posterior to bregma (Figure 5, green dots). These injections corresponded to the site targeted for our *in vitro* electrophysiological recordings. Three animals were excluded from drug-seeking analyses based on misplaced injection sites. In one case, both injectors were ventral to the orexin cell field (pictured in Figure 5; pink dots). This animal exhibited much higher reinstatement behavior following L-AP4 compared to the group average of animals that received accurate injections of L-AP4 (active lever presses=27, compared to a group L-AP4 average 15.91±2.98 SEM). Another animal had one accurate injection site in one hemisphere, but the other was located ventral to the orexin cell field (pictured in Figure 5; pink dots). In another, both injectors were located in the very rostral extent of the orexin cell field (−2.28mm; not pictured) rather than the center of the orexin cell field as will all other cases (Figure 5, green dots). For Fos and locomotor control experiments, all cannula placements were accurate; microinjections ranged from 2.64mm to 2.92mm posterior to bregma (Figure 5, blue dots).

**Figure 5.**
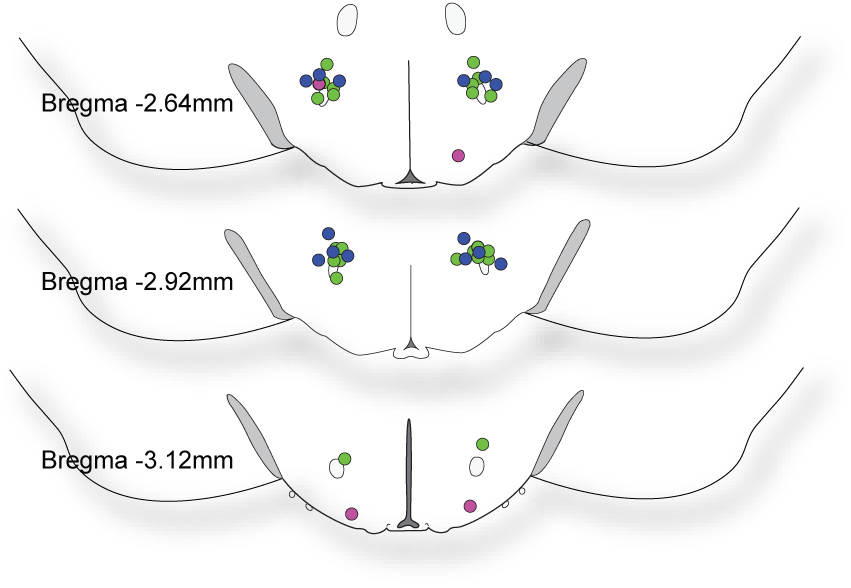
Localization of LHA-directed L-AP4 microinjections. Schematics of the hypothalamus showing accurately placed infusion sites for drug-seeking (green) and Fos/locomotor control experiments (blue). Misplaced infusions from the drug-seeking experiment that were excluded from behavioral analyses are depicted in pink.

### 3.2. Effects of i.p. cocaine on plasticity in LHA orexin cells and impact of GPIII mGluR activation

#### 3.2.1. Cocaine produced locomotor sensitization

Orexin-GFP mice that were exposed to cocaine exhibited a significant increase in locomotor activity compared to baseline (Figure 6A). Locomotor activity in response to cocaine was significantly higher in the final 3d of cocaine exposure (t(4)=6.976, *P*<0.05) compared to the first 3d of cocaine exposure (t(4)=3.539, *P*<0.05; Figure 6A), indicating behavioral sensitization (main effect of ‘treatment’, F_2, 4_=24.33, *P*=0.0058). In contrast, saline had no effect on locomotor activity (Figure 6B).

**Figure 6.**
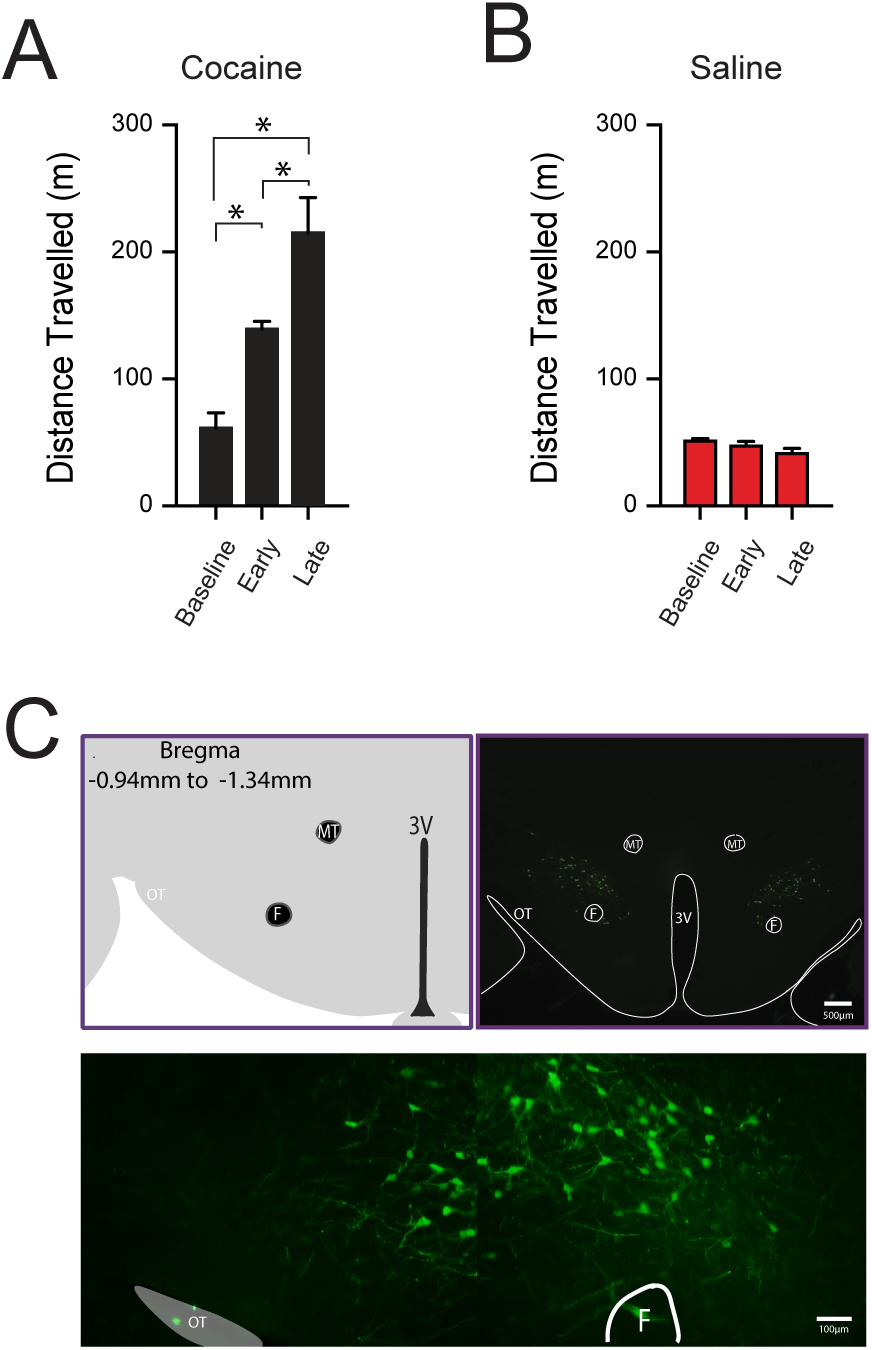
Cocaine induces locomotor sensitization & distribution of GFP-orexin cells in the LHA. *A*, repeated cocaine injections produced a significant increase in locomotor activity (distance travelled) as compared to saline-treated animals (^*^*P*<0.05, relative to baseline). Locomotor activity in response to cocaine was significantly higher in the final 3d of cocaine exposure compared to the first 3d of cocaine exposure (^*^*P*<0.05; Fig 1, left panel). *B*, Saline had no effect on locomotor activity. *C*, GFP expression is contained exclusively to orexin cells.

#### 3.2.2. Cocaine-induced changes to LHA-orexin circuits persist for 2 weeks post-cocaine

To determine whether cocaine-induced plasticity in orexin cell circuits persisted into withdrawal, we recorded orexin cells in hypothalamic slices taken from animals that had undergone 14d home cage abstinence following 7d of cocaine injections. Cocaine-exposed animals displayed a paired-pulse depression compared to controls (t_23_=2.598, *P*<0.05, cocaine vs saline, cell yield/animal = 5.5 vs 3.2 respectively, Figure 7A) and a significant increase in the AMPA:NMDA ratio in orexin cells from 14d withdrawn animals was also observed (t_13_=−2.731, *P*<0.05, cocaine vs saline, cell yield/animal = 3.0 vs 1.8 respectively, Figure 7B).

**Figure 7.**
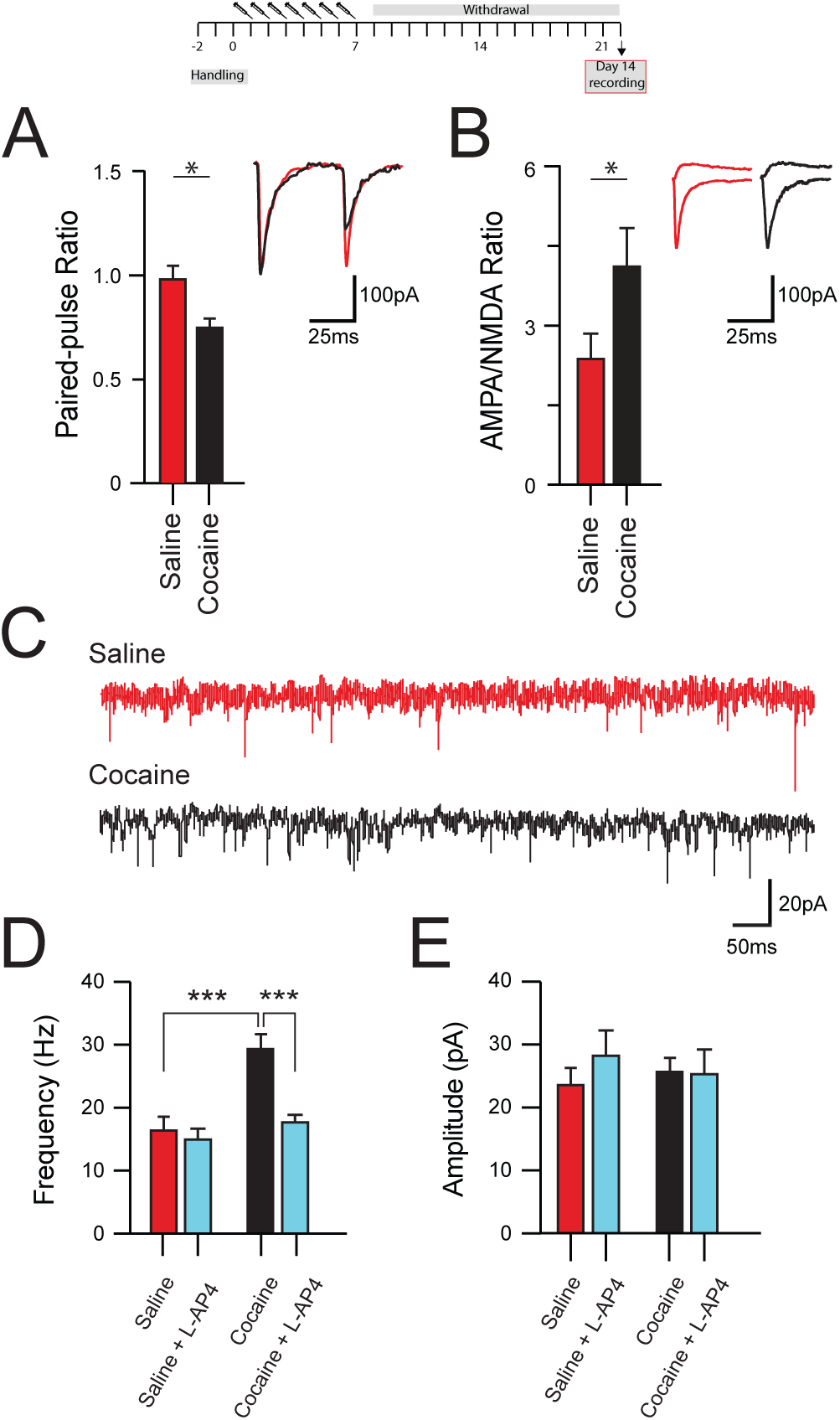
Cocaine-induced changes in paired-pulse ratio, AMPA:NMDA ratio, and mEPSCs frequency on D14 of cocaine withdrawal. *A*, bar graph compares group data indicating that recordings from cocaine-exposed mice exhibited paired-pulse depression (^*^*P*<0.05). Inset shows representative traces of eEPSCs recorded in orexin cells from saline- and cocaine-exposed animals. *B*, bar graph compares group data indicating that recordings from cocaine-exposed mice exhibited an increase in AMPA:NMDA ratio (^*^*P*<0.05). Inset shows representative overlaid AMPA-mediated (recorded at −70mV) and NMDA-mediated (recorded at +40mV, in 10μM CNQX) eEPSCs. *C*, traces show representative mEPSCs recordings of orexin cells from animals exposed to either saline (top) or cocaine (bottom). *D*, Bar graphs compare the mEPSCs frequency recorded in orexin cells from saline-exposed animals, saline + bath application of L-AP4, cocaine-exposed animals, and cocaine + bath application of L-AP4 at withdrawal day 14. Cocaine exposure significantly increased the mEPSCs frequency compared to saline control (^**^*P*<0.01), and application of L-AP4 reduces the mEPSCs frequency back to control level. *E*, bar graphs compare the mEPSCs amplitude between saline-, saline + LAP4, cocaine-, and cocaine + L-AP4. L-AP4 did not affect mEPSC amplitude in both treatment groups. The data presented are mean +/- SEM.

We also recorded mEPSCs properties in LHA cells 14d after final cocaine exposure (Figure 7C). mEPSCs frequency was significantly increased after 14d cocaine abstinence compared to saline (t_26_=-3.796, *P*<0.01, cocaine vs saline, cell yield/animal = 2.8 vs 3.5 respectively, Figure 7D), whereas amplitude (*P*>0.05, Figure 7E) and decay kinetics (*P*>0.05) were unchanged. Bath application of L-AP4 significantly reduced mEPSCs frequency in 14d cocaine withdrawn animals (t_19_=3.041, *P*<0.01, pre L-AP4 vs post L-AP4 treatment, cell yield/animal = 2.8 vs 2.3 respectively, Figure 7D), but had no effect on mEPSCs frequency in saline animals (*P*>0.05). L-AP4 had no effect on amplitude (*P*>0.05, Figure 7E) or decay kinetics in both groups (*P*>0.05).

## 4. DISCUSSION

We show that microinfusions of the group III mGluR agonist L-AP4 into the LHA orexin field effectively suppressed orexin cell activity and reduced cocaine-seeking behavior following 14d withdrawal in rats. Importantly, intra-LH L-AP4 had no apparent effects on general locomotor activity or self-administration of sucrose pellets. We confirm that L-AP4 acts via a presynaptic mechanism by showing that withdrawal-induced increases in mEPSCs frequency are reversed by L-AP4 in recordings of identified orexin cells in mice. Together, these findings indicate that agonism of group III mGluRs can suppresses presynaptic plasticity onto orexin cells following cocaine withdrawal. Direct infusion of L-AP4 into the LHA reduces drug-seeking behavior following abstinence, thus identifying this system as a possible target to normalize orexin cell activity following cocaine use.

### 4.1. Agonism of LHA group III mGluRs reduces orexin cell activity and reduces cocaine seeking behavior

Agonism of group III mGluRs with L-AP4 has previously been shown to inhibit synaptic input onto orexin cells in slice (Acuna-Goycolea *et al.*, 2004). Here, we extend these findings to show that infusions of L-AP4 directly into the LHA orexin cell field reduced activity of orexin cells following 24h food deprivation. We chose this stimulus based on previous studies that demonstrated food deprivation increases presynaptic input onto orexin cells, as measured by mEPSCs frequency and the number of vGlut2+ve inputs in close apposition to orexin cells, as well as increased Fos expression in orexin cells (Mieda *et al.*, 2004; Horvath & Gao, 2005). We show that L-AP4 infusions reduced the amount of Fos by approximately half; this finding aligns well with electrophysiological evidence that bath application of L-AP4 inhibits spontaneous excitatory activity in orexin cells by roughly the same magnitude (Acuna-Goycolea *et al.*, 2004). It is important to point out however, that L-AP4 microinfusions likely also affected the activity of non-orexin cells in this region, especially given the large heterogeneity of cell populations in the LH (Bonnavion *et al.*, 2016). Nevertheless, these experiments identify the group III mGluR system as a potential target to suppress the activity of orexin cells in behaving animals.

Previous studies have shown that orexin cells are activated by cocaine-associated stimuli (Martin-Fardon *et al.*, 2018), and that their signaling is critical for the expression of drug-seeking behavior elicited by these stimuli (Martin-Fardon & Weiss, 2014). Here, we show that infusions of L-AP4 directly into the LHA orexin cell field significantly reduced drug-seeking elicited by reintroduction of cocaine-paired discriminative-stimuli following 14d withdrawal in rats previously trained to self-administer cocaine. Although we did not directly assess the effect of L-AP4 on orexin cell activity in this test, our findings in food-deprived rats, as well as previous electrophysiological evidence (Acuna-Goycolea *et al.*, 2004), indicates that this behavioral effect was mediated by a suppression of orexin cell activity elicited by the drug-context. This finding aligns well with previous studies that have reported reduced stimulus-driven cocaine-seeking following abstinence or extinction in orexin knock-out mice (Steiner *et al.*, 2018) or following either systemic or local pretreatment with an orexin-1 receptor antagonist (Smith *et al.*, 2010; James *et al.*, 2011; Mahler *et al.*, 2013). While this is the first study to directly implicate LHA group III mGluRs in drug-seeking, these same receptors have been shown to regulate other forms of drug behavior in caudate. Intra-caudate microinjections of L-AP4 blocked hyperlocomotion induced by an acute systemic injection of cocaine, amphetamine or apomorphine, and this effect was blocked by co-administration of the group III *antagonist*, a-methyl-4-phosphonophenylglycine (MPPG; (Mao & Wang, 2000). Thus, group III mGluRs may act at various sites to regulate several glutamate-dependent drug behaviors, potentially increasing the utility of compounds targeting these receptors in blocking addiction-like behavior.

Importantly, L-AP4 had no effect on lever pressing for sucrose pellets or spontaneous locomotor activity in an open field. These findings indicate that L-AP4 did not affect general locomotor activity, a finding that is consistent with a previous study showing that bilateral intra-caudate microinjections of L-AP4 at a similar dose to that used here had no effect on basal locomotor activity (Mao & Wang, 2000). As such, the effects of L-AP4 on cocaine-seeking behavior are likely related to motivation, rather than impaired motor behavior. Importantly however, it is unclear whether group III mGluRs uniquely mediate motivated drug-seeking; indeed, we did not test the effect of LH microinjections of L-AP4 on motivated food-seeking (such as on a reinstatement or progressive ratio paradigm). Given the known role for orexin in mediating motivated food behavior (Cason & Aston-Jones, 2013b; a), it might be expected that similar findings would be observed in such tests, although future studies will be needed to test this hypothesis.

Interestingly, intra-LH microinjections of L-AP4 increased the amount of time rats spent in the centre square of the open field apparatus during locomotor testing, indicative of an anxiolytic effect. This observation aligns with several studies implicating orexin signaling in the expression of anxiety behavior; exogenous orexin administration of orexin peptides (Ida *et al.*, 1999; Ida *et al.*, 2000) or optogenetic stimulation of orexin cells (Heydendael *et al.*, 2013; Bonnavion *et al.*, 2015) promotes stress behaviors, whereas orexin receptor antagonists generally suppress stress- and anxiety-like behaviors (Plaza-Zabala *et al.*, 2010; Staples & Cornish, 2014; Vanderhaven *et al.*, 2015; James *et al.*, 2017a). Some evidence indicates that the medial orexin cell populations (DMH and PF) specifically modulate anxiety and stress-behavior, whereas the lateral population plays a stronger role in regulating reward behavior (see (James *et al.*, 2017b) for review). Here, injections of L-AP4 were made into the bulk of the orexin cell field (immediately dorsal to the fornix) and thus both medial and lateral orexin cell populations were affected, perhaps accounting for the effects of this manipulation on both reward and anxiety behaviors.

### 4.2. Group III mGluR agonism reverses presynaptic plasticity within LHA orexin circuits following cocaine withdrawal

We next sought to characterize, at the synaptic level, the mechanisms through which L-AP4 acts to suppress orexin cell activity. To do this, we used a well-validated mouse model in which GFP is driven by the human orexin promoter, thus allowing us to record from identified orexin cells in slice. We exposed mice to 7d of experimenter-administered cocaine, a regimen that we have previously shown to produce plasticity within LHA orexin circuit that is identical to that produced by cocaine self-administration (Yeoh *et al.*, 2012). This regimen was associated with a prototypical sensitization of locomotor activity, thought to reflect neuroadaptive changes governing the development of drug-dependence (Cornish & Kalivas, 2001). We have previously reported enhanced excitatory input onto orexin cells 1d after cocaine sensitization or self-administration; here we extend these findings to show a significant paired-pulse depression at excitatory synapses, accompanied by an increase in mEPSCs frequency, on orexin-GFP cells following 14d withdrawal from a cocaine sensitization regime. This finding suggests that the increased drive to orexin cells persists well after cessation of drug exposure, potentially contributing to the persistent risk of relapse observed in animals and humans. We did not observe a change in mEPSCs amplitude, suggesting a lack of postsynaptic plasticity at orexin cells in response to cocaine. This finding is surprising given our observation using electrically evoked ESPCs that cocaine exposure was associated with a significant increase in AMPA:NMDA ratio. Importantly however, mEPSCs effectively samples all excitatory inputs on orexin cells, providing an average of the excitatory drive from all afferent pathways (local and extrinsic). In contrast, electrically evoked EPSCs, assess a subset of excitatory inputs determined by the placement of our stimulating electrode, which was within the LHA (local). Thus, a more restricted LHA-based population of inputs may be potentiated by cocaine and revealed only after local stimulation. Future studies could address this issue using virally-mediated stimulation of specific afferents.

Our findings are largely consistent with a previous study that reported that repeated i.p. injections of cocaine, but not a single injection, increased the AMPA:NMDA ratio and was associated with enhanced mEPSCs amplitude in GFP-orexin cells (Rao *et al.*, 2013). This study also showed that cocaine facilitated the induction of LTP by high frequency stimulation in orexin cells. Interestingly however, this study reported that these measures of plasticity persisted for only 5d; a significantly shorter period of time than that reported here (14d). While the reasons for these differences is unclear, it is possible that the higher dose of cocaine (15 versus 10mg/kg) and the duration of cocaine injections (7d versus 3d) used in the current study contributed to the more persistent effects reported here. Regardless, both studies point to significant rewiring of LHA orexin system following repeated cocaine exposure; it is possible that these changes augment the extent to which orexin cells are recruited by other salient stimuli, such as stress or cues linked with drug-taking, even after significant periods of abstinence, and thus may underlie the persistent relapsing nature of addiction .

Consistent with our *in vivo* data, we show that activation of group III mGluRs with L-AP4 significantly reduced mEPSCs frequency in orexin-GFP cells recorded from cocaine-exposed animals. L-AP4 had no effect on mEPSC amplitude, indicating a selective normalization of presynaptic plasticity in these animals. Interestingly, L-AP4 had no effect on either mEPSCs frequency or amplitude in saline-treated animals. This indicates that cocaine produced a persistent change in the expression or function of LHA mGluRs, as has been reported in other brain regions under different circumstances (Gordon & Bains, 2003). Future studies will need to explore this possibility by probing the upstream signaling pathways (e.g. PKC) responsible for regulating GP III mGluR function.

Our finding that L-AP4 did not affect mEPSCs frequency in saline-treated rats contrasts with those of Acuna-Goycolea *et al* (2004), who reported a reduction in spontaneous glutamate- and GABA-mediated synaptic currents following bath applications of L-AP4 in drug-naïve animals. Although speculative, it is possible that repeated saline injections in our study resulted in a shift in baseline mEPSCs frequencies or altered mGluR receptor density. It should also be highlighted that L-AP4 is thought to be more selective for mGluR4 or 8 receptors at the concentration used in our experiments, indicating that the group III mGluR responsible for these effects is likely through the activation of one or both of these ‘high-affinity’ subtype (Gereau & Swanson, 2008). Further studies are required to delineate the specific subtype involved.

One important caveat of our study is that our behavioral experiments were carried out in self-administering rats whereas the electrophysiological experiments were conducted in experimenter-exposed mice. The discriminative stimulus operant drug-seeking model used here has been characterized far more extensively in rat compared to mouse (Dayas *et al.*, 2008; Martin-Fardon & Weiss, 2013; 2014; McGlinchey & Aston-Jones, 2017; Martin-Fardon *et al.*, 2018); thus our behavioral data can be more readily compared with previous studies. In contrast, the orexin-GFP transgenic mouse line has been verified and characterized across several studies, whereas comparable studies in rat are limited (Yamanaka *et al.*, 2003a; Yamanaka *et al.*, 2003b; Sakurai *et al.*, 2005). Therefore, some caution should be taken when considering how our electrophysiological findings relate to the reported behavioral data, and vice versa. However, we note that all published literature to date points to a large degree of overlap in the orexin system between rat and mouse in terms of topography (de Lecea *et al.*, 1998; Sakurai *et al.*, 1998; Sakurai *et al.*, 1999; Stricker-Krongrad *et al.*, 2002), projections and receptor distribution (Peyron *et al.*, 1998; Chen *et al.*, 1999; Marcus *et al.*, 2001; Lin *et al.*, 2002; Puskás *et al.*, 2010; Ch’ng & Lawrence, 2015), electrophysiological properties (Yeoh *et al.*, 2012) and role in reward-seeking behaviors (James *et al.*, 2017b; Schmeichel *et al.*, 2018; Steiner *et al.*, 2018). Moreover, we have previously reported that plasticity at orexin cells occurs similarly between rats that self-administer cocaine and mice that receive experimenter-administered cocaine (as was the case here) on d1 of withdrawal (Yeoh *et al.*, 2012). Nevertheless, future studies should confirm that the plasticity that we observed on withdrawal d14 in mice is similar in rats with a history of self-administration.

### 4.3. Conclusions

In conclusion, we show that agonism of LHA group III mGluRs inhibits orexin cell activity and drug-seeking behaviour *in vivo*, and reverses increases in presynaptic plasticity at orexin cells *in vitro*. The exact source of increased excitatory drive to LH circuits remains to be determined, with possible roles for both intra- and extra-hypothalamic sources; it will be important for future studies to examine these possibilities using circuit-specific manipulations. Regardless, the present findings highlight GPIII mGluRs as a potential novel target for pharmacotherapies designed to suppress hyperactivity of orexin cells during abstinence and ameliorate relapse risk.

